# Di-arginine and FFAT-like motifs retain a subpopulation of PRA1 at ER-mitochondria membrane contact sites

**DOI:** 10.1101/2020.01.27.922260

**Authors:** Ameair Abu Irqeba, Judith Mosinger Ogilvie

## Abstract

Prenylated Rab Acceptor 1 (PRA1/Rabac1) is a four-pass transmembrane protein that has been found to localize to the Golgi and promiscuously associate with a diverse array of Rab GTPases. We have previously identified PRA1 to be among the earliest significantly down-regulated genes in the *rd1* mouse model of retinitis pigmentosa, a retinal degenerative disease. Here, we show that an endogenous subpopulation of PRA1 resides within the endoplasmic reticulum (ER) at ER-mitochondria membrane contact sites in cultured mammalian cells. We also demonstrate that PRA1 contains two previously unidentified ER retention/retrieval amino acid sequences on its cytosolic N-terminal region: a membrane distal di-arginine motif and a novel membrane proximal FFAT-like motif. Using a truncation construct that lacks complete Golgi targeting information, we show that mutation of either motif leads to an increase in cell surface localization, while mutation of both motifs exhibits an additive effect. We also present evidence that illustrates that N- or C- terminal addition of a tag to full-length PRA1 leads to differential localization to either the Golgi or reticular ER, phenotypes that do not completely mirror endogenous protein localization. The presence of multiple ER retention motifs on the PRA1 N-terminal region further suggests that it has a functional role within the ER.

## INTRODUCTION

The *rd1* mouse is an animal model of retinitis pigmentosa, a disease that affects nearly one in four thousand, leading to degeneration of photoreceptors and eventual blindness of affected individuals (Farber et al., 1994; Hartong et al., 2006). We have previously identified Prenylated Rab Acceptor 1 (PRA1) as a gene that is significantly down-regulated at postnatal day 2 (P2) in the *rd1* mouse retina (Dickison et al., 2012), particularly notable since cell death begins around P10. We have also shown that PRA1 is enriched in normal photoreceptors, is not trafficked to the outer segment, and is mis-localized in *rd1* photoreceptors prior to the onset of degeneration (Dickison et al., 2012).

PRA1 is highly conserved in eukaryotes and ubiquitously expressed in mammalian tissues (Bucci et al., 2001). Previous work has suggested that it plays an important role in protein trafficking, specifically in recycling Rab GTPases back to donor membranes (Hutt et al., 2000; Sivars et al., 2003). Rabs are members of the Ras superfamily that regulate vesicular targeting and trafficking within eukaryotic cells. The recycling of Rabs back to their donor membrane requires a GDP dissociation inhibitor (GDI) to mask the prenyl tail from the aqueous environment after extraction from the target membrane. Rabs are known to strongly associate with GDIs to form a complex, which has led some to postulate that there must be an effector, a GDI displacement factor (GDF), on the donor membrane that participates in the dissociation process and re-insertion of the Rab prenyl tail into the donor membrane (Oesterlin et al., 2012; Ohya et al., 2009; Pfeffer and Aivazian, 2004; Schalk et al., 1996). The only proposed eukaryotic Rab GDF thus far is PRA1 (Abdul-Ghani et al., 2001; Liang and Li, 2000).

PRA1 promiscuously associates with many Rabs, although it seems to have a preference for those that are endosomally localized (Bucci et al., 1999; Calero and Collins, 2002; Janoueix-Lerosey et al., 1995; Martincic et al., 1997; Ohya et al., 2009; Sivars et al., 2003). While previous studies have alluded to a role for PRA1 in Rab trafficking, in-vivo evidence that supports this proposed function has been lacking (Ohya et al., 2009; Sivars et al., 2003). In fact, knock-out of the PRA1 homolog in yeast demonstrates that it is non-essential, with no evident phenotype in the endosomal system and no change in Rab GTPase localization (Cabrera and Ungermann, 2013; Geng et al., 2005). Furthermore, in-vivo evidence points to a more structural role within the early secretory pathway as knock-down or knock-out of PRA1 leads to abnormal ER and Golgi phenotypes (Geng et al., 2005; Lee et al., 2017; Liu et al., 2011; Simpson et al., 2012). We recently used an unbiased approach that takes into account the membrane association of PRA1 to screen for novel binding partners and failed to identify any Rab GTPases (Abu Irqeba and Ogilvie, 2019). Together, this suggests that PRA1 may have other roles that have not been elucidated.

To gain insight into the functional role that PRA1 plays in photoreceptors and mammalian cells in general, we studied the localization of endogenous PRA1 and the minimal trafficking motifs responsible for its retention within the early secretory pathway. Previously, others have demonstrated that PRA1 is a Golgi resident (Abdul-Ghani et al., 2001; Liang and Li, 2000). Here, we show that endogenous PRA1 does not strictly localize to the Golgi. In NIH3T3 cells, a large population also resides at ER-mitochondria membrane contact sites. We present evidence that a novel, but highly conserved FFAT-like motif and a di-arginine ER retrieval motif on the PRA1 cytosolic N-terminal region each play a role in the retention of a subpopulation within the ER. Mutation of these motifs in PRA1 constructs with incomplete Golgi targeting information leads to an increase in cell surface localization within epithelial cells. We further demonstrate that the addition of tags to the full length PRA1 open reading frame obfuscates its true endogenous localization. To our knowledge, this is the first time that endogenous mammalian PRA1 has been localized to the ER.

## METHODS

### Animals

C57Bl/6J mice were used for all experiments. Animals were housed in 12hr/12hr light/dark conditions with food and water ad lib. All experiments conformed to the National Institutes of Health Guidelines on Laboratory Animal Welfare using procedures that were approved by the Saint Louis University Institutional Animal Care and Use Committee.

### Plasmids and cloning

All cloning reagents were purchased from New England Biolabs (Ipswich, MA, USA). All oligonucleotides used were ordered from Integrated DNA Technologies (Coralville, IA, USA).

See Table S1 for primer sequences. The pCAGIG vector was made available by Connie Cepko via Addgene (Cambridge, MA, USA; plasmid #11159) (Matsuda and Cepko, 2004). To generate the mCherry-sec61b construct, mCherry was amplified from Addgene plasmid #49155 with primers containing an EcoRI site upstream and Kpni, MluI, and BglII sites downstream of the PCR product and subsequently cloned into the EcoRI/BglII sites on the pCAGIG vector. Sec61b was amplified from mouse retinal cDNA and cloned downstream of mCherry using the KpnI/MluI restriction enzyme sites. To generate mMannII-mCherry, mCherry was cloned within the pCAGIG vector EcoRI/BglII sites after amplification with a forward primer that has an EcoRI and MluI restriction enzyme sites and reverse primer with a BglII restriction enzyme site. Mouse Mannosidase II amino acids 1-116 was amplified from Addgene plasmid #65261 using a forward primer that contains an XbaI site and a reverse primer with an MluI site and further subcloned upstream of the mCherry open reading frame.

Overlap-extension PCR was used to generate the EGFP^A206K^ mutant that was fused N/C-terminally to all PRA1-GFP/GFP-PRA1 constructs. Mouse PRA1 cDNA was obtained from Origene (Rockville, MD, USA; Cat# MC200290). The PRA1-GFP construct was made by amplification and cloning of the PRA1 ORF into the pCAGIG vector using the XhoI/MscI restriction sites. To generate the GFP-PRA1 construct, EGFP^A206K^ was amplified with a forward primer that contains an XhoI site and a reverse primer that contains both MluI and BglII sites and cloned into the PCAGIG vector using the XhoI/BglII sites. PRA1 was then subcloned downstream of GFP using the MluI and BglII sites.

For the cell surface assay, the Snorkel-Tag open reading frame was re-constructed using overlapping oligos with a second FLAG tag added to its C-terminus to increase anti-FLAG antigenicity (Brown et al., 2013). The Snorkel-Tag was cloned into the XhoI and BglII sites on the pCAGIG vector with a KpnI site added between the XhoI site and the Snorkel-Tag open reading frame. PRA1 constructs were then subcloned into the XhoI/KpnI sites in-frame with the downstream Snorkel-Tag. All PRA1 mutations described in this manuscript were introduced via overlap-extension PCR (Ho et al., 1989).

All constructs used in this study were verified via Sanger sequencing at either the Washington University School of Medicine Protein and Nucleic Acid Chemistry Laboratory (St. Louis, MO, USA) or Eurofins Sequencing (Louisville, KY, USA).

### Cell culture

COS-7 cells [ATCC CRL-1651] and NIH3T3-L1 cells [ATCC CL-173] were acquired from American Type Culture Collection (ATCC, Manassas, VA). Cell lines were validated via morphology and growth curve analysis prior to use. Cells were cultured in DMEM with 10% fetal bovine serum (Sigma-Aldrich, St. Louis, MO) and 1X antibiotic/antimycotic solution (Gibco, Waltham, MA). Transfections were carried out using Lipofectamine 2000 (ThermoFisher Scientific, Waltham, MA) in 12 and 24 well plates according to manufacturer’s instructions.

### Immunochemical staining

Eyecups were harvested at P21, fixed in 4% paraformaldehyde, washed in 0.1 M phosphate buffer, cryoprotected in 30% sucrose overnight, and embedded in OCT (Sakura, Torrance, CA, USA), as described previously (Dickison et al., 2012). Eyecups were cut into twelve-micron sections using a Leica CM 1850 cryostat. Cell cultures were grown on coverslips. For labeling with the 3F3A antibody, cell cultures were fixed using −80°C super-cooled 80/20 % v/v methanol/acetone mixture and then placed at −20°C for 20 minutes. Cells were then rehydrated with a PBS rinse every 10 minutes for 60 minutes total. For all other cell culture labeling, a standard 4% paraformaldehyde fixation protocol was used. No immunofluorescence was observed in control experiments omitting primary antibodies.

The following antibodies (diluted in blocking solution) were used in this study: rabbit anti-PRA1 (1:100, Abgent, San Diego, CA, Cat #AP9049a; polyclonal generated against amino acids 1-30; validated in Fig. S1), rabbit anti-PRA1 (1:100, Proteintech, Rosemont, IL, Cat #10542-1-AP; polyclonal generated against full length protein; validated in Fig. S2), mouse anti-GM130 (1:1000, BD Biosciences, San Jose, CA, Cat #610822), mouse anti-KDEL (1:200, Abcam, Cambridge, UK, Cat #ab12223), goat anti-Calnexin (1:50, Santa Cruz, Dallas, TX, Cat #sc- 6465), mouse anti-Ctbp2/Ribeye (1:100, BD Biosciences, San Jose, CA, Cat #612044), and rat anti-Gp78/AMFR (1:50, EMD Millipore, St. Louis, MO, Cat #MABC949). The mitochondria were labeled with MitoTracker DeepRed FM according to manufacturer’s instructions (Cat #M22426 ThermoFisher Scientific). All secondary antibodies used were Alexa Fluor (488/555/647) dyes from ThermoFisher Scientific.

For retinal sections, after the application of secondary antibodies, the nucleus was stained with TO-PRO-3 iodide (1:500 in phosphate buffered saline (PBS), ThermoFisher Scientific Cat #T3605) for 15 minutes. Sections were then washed in PBS for 15 minutes. Cell cultures were stained according to previously described procedures (Harlow and Lane, 2006). Vectashield mounting medium with or without DAPI (VWR, Radnor, PA) was applied to each retinal section or cultured cells prior to the application of coverslips.

### Confocal microscopy and data analysis

Retinal images were acquired using a Zeiss LSM 510 Meta Pascal confocal microscope (63x oil objective). All cultured cell line images were acquired using an Olympus FV1000 confocal microscope (60x oil objective). All settings were used consistently for experiments stained with the same antibodies. Image processing was completed using ImageJ software (National Institutes of Health). All immunohistochemical data is representative of three non-adjacent sections from each of four different retinas or, for cell cultures, of four different transfections in four separate wells.

### Cell Surface Assay

The cell surface assay protocol was carried out in 24 well plates. COS-7 cells were plated at 10^4^ cells per well 24 hours prior to transfection. PRA1^1-131^-Snorkel-Tag fusions were used to quantify ER retention efficacy. The protocol for colorimetric quantification of cell surface protein localization was carried out as described previously (Andrisse et al., 2013; Ishikura et al., g of each construct was mixed with 0.6 μl of Lipofectamine 2000 in Opti-MEM, incubated for 30 minutes, then added to cells. Six hours after transfection, fresh media was added to each well. Twenty-four hours after transfection, cells were washed twice with room temperature PBS and new media was added. Forty-eight hours after transfection, cells were washed with PBS. A blocking solution (1% milk in PBS) was then added to cells for 20 minutes and subsequently replaced with an anti-FLAG antibody (1:1000 in blocking solution) (Cat #F3165, Sigma-Aldrich) for one hour. Cells were then washed 8X with PBS, fixed for 10 minutes with 4% paraformaldehyde, washed once with PBS, and the fixative was quenched with 1 mM glycine in PBS for 10 minutes. Cells were blocked for 15 minutes in blocking solution followed by incubation in an anti-mouse HRP secondary antibody diluted in blocking solution (1:750) for one hour. Cells were then washed 6X with PBS. PBS was then removed and OPD assay solution (Cat # P9187, Sigma-Aldrich) was added for 30 minutes in the dark, the reaction was then halted using 125 μl of 3 N HCl. Absorbance was read at 492nm using a Bio-Tek Synergy H1 plate reader (n=9 for each construct). GFP transfected and non-transfected wells were used as a control. No difference in background signal between the two controls was observed. The GFP open reading frame was expressed using the same vector backbone as the Snorkel-Tag fusion constructs.

### Statistical Analysis

After all absorbance data was averaged for each condition, the average for non-transfection control wells was subtracted from all readings to remove background. In order to determine the relative fold-change of each construct compared to the wild-type, the resulting averages were divided by the average for the wild-type construct. Statistical significance was determined using one-way ANOVA followed by a least significant difference (LSD) Post-Hoc test. A p value < 0.05 was considered significant.

## RESULTS

### Endogenous PRA1 does not strictly localize to the Golgi

We have previously identified PRA1 as a gene that is significantly down-regulated prior to the onset of photoreceptor degeneration in the *rd1* retina (Dickison et al., 2012). We sought to replicate previous results that established PRA1 as a Golgi resident (Abdul-Ghani et al., 2001; Liang and Li, 2000). PRA1 is known to be a multi-pass transmembrane protein (Fig. 1A) (Lin et al., 2001). Consistent with our previous report, endogenous PRA1 labeling in photoreceptors confirmed some co-localization with the Golgi marker GM130 (Fig. 1B) (Dickison et al., 2012; Nakamura et al., 1995). We also found that PRA1 partially co-localizes with the photoreceptor synapse marker Ribeye (Fig. 1C) (Fenster et al., 2000). In photoreceptors, the Golgi lies within the inner segment distal to the outer nuclear layer; yet staining for PRA1 demonstrates a localization pattern spread throughout the cell (Fig 1B). The distinct morphology and small diameter of mouse photoreceptors makes it difficult to discern among the subcellular compartments. To better characterize the subcellular localization, we turned to cultured NIH3T3 cells.

**Fig. 1.**
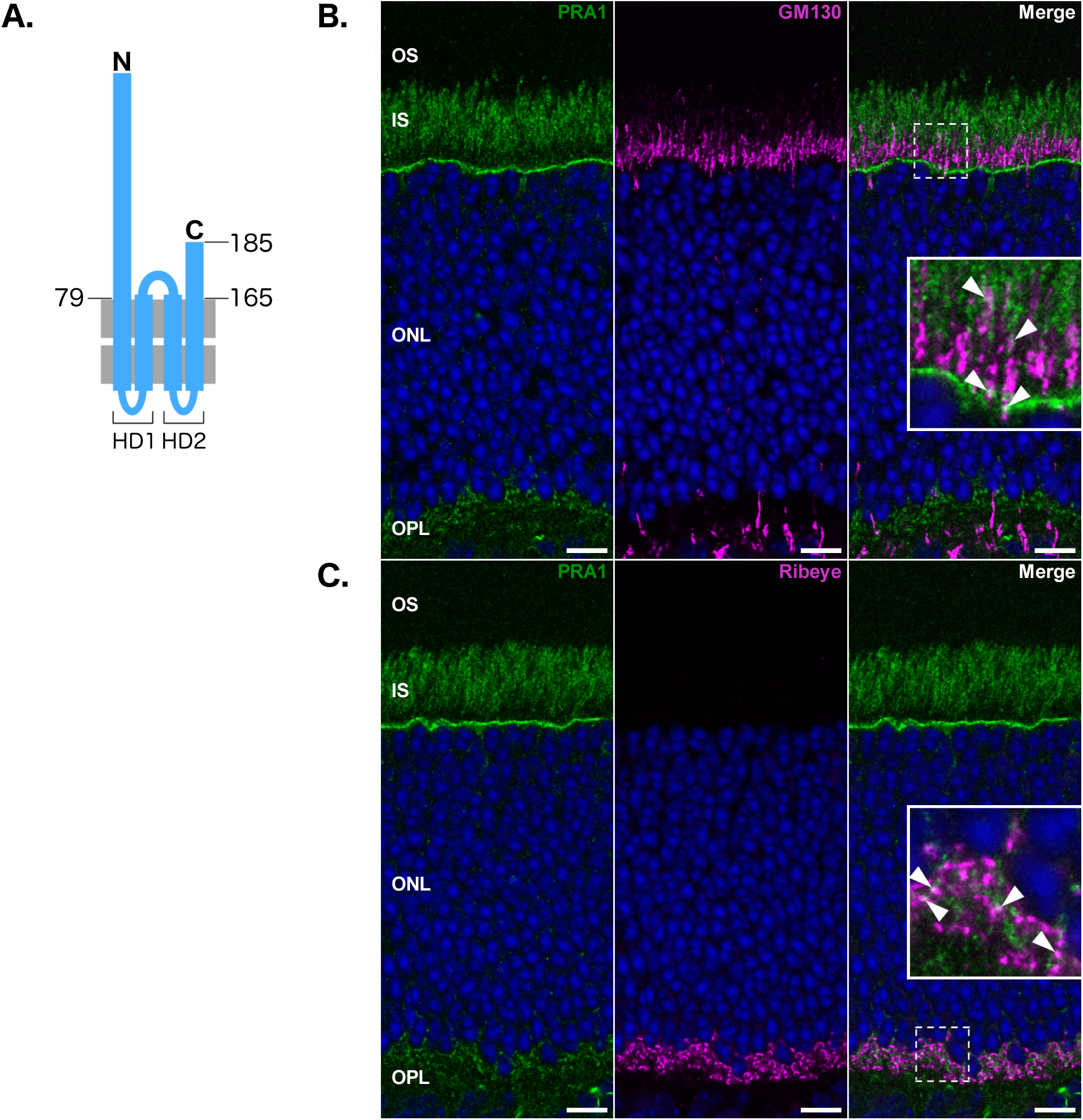
Endogenous PRA1 does not strictly localize to the Golgi in mature photoreceptor cells. (A) Schematic showing PRA1 topology. (B) PRA1 partially co-localizes with both the Golgi marker GM130, and the ribbon synapse marker Ribeye (C) as indicated by solid arrowheads. Images are representative of three non-adjacent sections from four retinas and are compiled from a single 1 μm z-section. Insets are five-fold magnifications of an area within the displayed images, highlighted with a dashed-line. Scale bar: 10 μm Green: PRA1, Magenta: organelle markers, Blue: nuclear labeling via TO-PRO-3, White: co-localization, OS: outer segment, IS: inner segment, ONL: outer nuclear layer, OPL: outer plexiform layer.

Consistent with observations in photoreceptors, endogenous PRA1 partially co-localizes with the Golgi marker GM130, but much of the protein lies outside of this organelle in NIH3T3 cells (Fig. 2A). Previous studies have localized PRA1 to the ER in both *Saccharomyces cerevisiae* and *Arabidopsis thaliana* (Alvim Kamei et al., 2008; Geng et al., 2005). In addition, PRA1 mRNA has been reported to co-fractionate with the ER in yeast, suggesting that the protein is translated across the ER membrane (Kraut-Cohen et al., 2013). We stained cells with both KDEL and Calnexin specific antibodies, and found that although some PRA1 does co-localize with these ER markers, much of it does not (Fig. 2B-C). We observed that PRA1 staining is characterized by a “string-like” pattern that radiates throughout the cell, which is reminiscent of mitochondrial localization. A specific and structurally distinct sub-region of the smooth ER is known to make contact with the mitochondria. This region of the ER only partially co-localizes with the reticular ER, and is distinct in its close apposition to the mitochondria (Goetz et al., 2007). Double labeling with PRA1 and MitoTracker, a mitochondrial marker, shows that PRA1 co-localizes extensively with mitochondria in NIH3T3 cells (Fig. 2D). This co-localization is not one-to-one in nature; it is evident that flanking regions of PRA1 immunoreactivity within these string-like domains does not completely lie at the interface with the mitochondria. This phenotype was confirmed using a second commercially available polyclonal antibody made against the full length protein (Proteintech), which also shows that PRA1 co-localizes with mitochondria in NIH3T3 cells (Fig. S2).

**Fig. 2.**
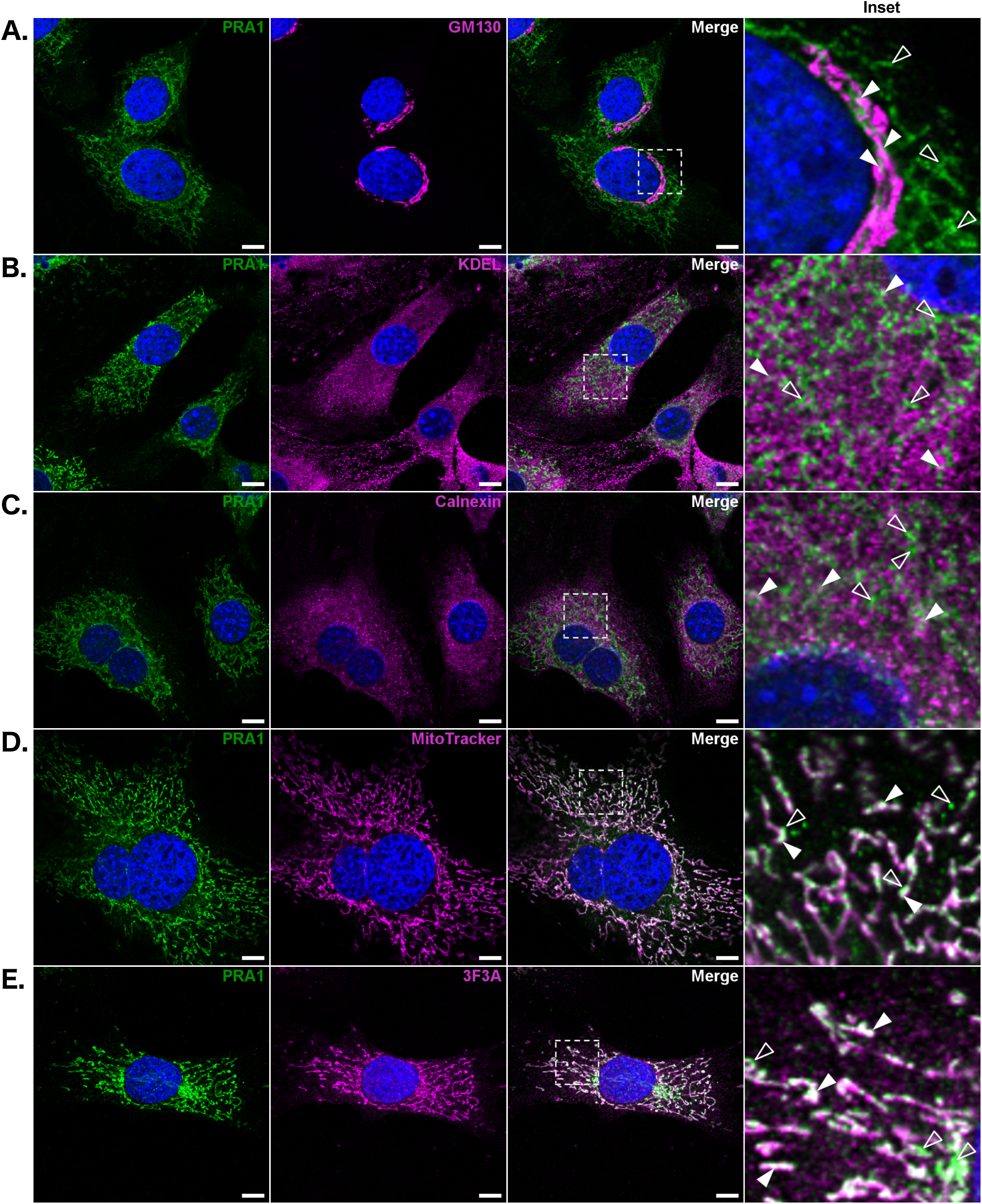
A subpopulation of endogenous PRA1 is localized to Gp78/AMFR positive ER-mitochondria membrane contact sites in NIH3T3 cells. PRA1 was co-stained with (A) GM130, (B) KDEL, (C) Calnexin, (D) MitoTracker, and (E) Gp78/AMFR (3F3A antibody). Solid arrowheads denote regions of co-localization; empty arrowheads denote regions of independent PRA1 localization. Images are representative of at least three acquisitions and are compiled from a 1 μm z-section. Insets are magnifications of an area within the displayed images, highlighted with a dashed-line. Scale bar: 10 μm Green: PRA1, Magenta: organelle specific marker, Blue: nuclear labeling via DAPI, White: co-localization.

The 3F3A antibody to Gp78/AMFR has previously been found to label a subpopulation of this protein that is localized to the smooth ER tubules that lie at the interface with the mitochondria (Goetz et al., 2007; Nabi et al., 1990). Co-localization of Gp78/AMFR (3F3A) and PRA1 further confirms that a subpopulation of PRA1 resides at ER-mitochondria membrane contact sites (Fig. 2E). Together, this data demonstrates that endogenous PRA1 localization extends beyond the Golgi in both photoreceptors and NIH3T3 cells. Specifically, within NIH3T3 cells, a sub-population of endogenous PRA1 localizes to ER-mitochondria membrane contact sites.

### Mutation of di-arginine and FFAT-like motifs on the PRA1 cytosolic N-terminal region leads to a significant increase in cell surface localization of constructs with incomplete Golgi targeting information

The topology of PRA1 has previously been characterized using in-depth biochemical studies (Fig. 1A) (Lin et al., 2001). The four-pass transmembrane structure is interrupted by an 18 amino acid central cytosolic stretch. Both N- and C-terminal regions are cytosolic, 78 and 20 amino acids in length, respectively. Two highly hydrophobic domains facilitate the association of PRA1 with the membrane: HD1 and HD2, each made up of two predicted membrane-spanning alpha helices.

Previously, it has been demonstrated that the short cytosolic C-terminal region contains a di-acidic motif that facilitates ER exit and delivery to the Golgi (Abdul-Ghani et al., 2001; Hutt et al., 2000; Liang and Li, 2000; Otte et al., 2001). Furthermore, the C-terminal valine is known to be important for dimerization of PRA1, which is also required for ER exit (Liang et al., 2004). Deletion of the cytosolic C-terminal region leads to ER retention, suggesting that the ER localization we observed is mediated by amino acid sequence information on the 78 amino acid long cytosolic N-terminal region, which has not been extensively studied (Liang and Li, 2000).

To screen for the possible existence of ER retention motifs on the PRA1 N-terminal domain, we used an ELISA-based cell surface assay approach. Targeted mutation of amino acid sequences responsible for retention to the ER should lead to an increase in the cell surface localization of PRA1 truncation constructs, as these mutants will be pushed out to the plasma membrane at a higher rate. To acquire an epitope that becomes extracellular and detectable upon PRA1 cell surface localization, we fused the PRA1^1-131^ construct to the Snorkel-Tag (Fig. 3C), a single pass transmembrane protein designed as a tool to survey cell surface localization for membrane proteins that have a cytosolic C-terminus (Brown et al., 2013). We specifically used the PRA1^1-131^ construct because it does not have the required Golgi targeting sequences that lie on the cytosolic C-terminal domain; furthermore, this truncation construct has previously been shown to reside within the ER by default (Liang and Li, 2000). We have also found that regions within both PRA1 transmembrane domains play an important role in facilitating Golgi delivery and retention (unpublished data). By omitting the HD2 domain and the cytosolic C-terminal region, the PRA1^1-131^ truncation construct inherently disrupts this Golgi retention mechanism, which allows for more efficient passage through the secretory pathway to the cell surface without changing the topology of the HD1 domain or modifying cytosolic N-terminal region trafficking motif distance from the membrane. Thus, this de-coupling strategy further optimizes the cell surface assay and allows for better output resolution without introducing other confounding variables.

**Fig. 3.**
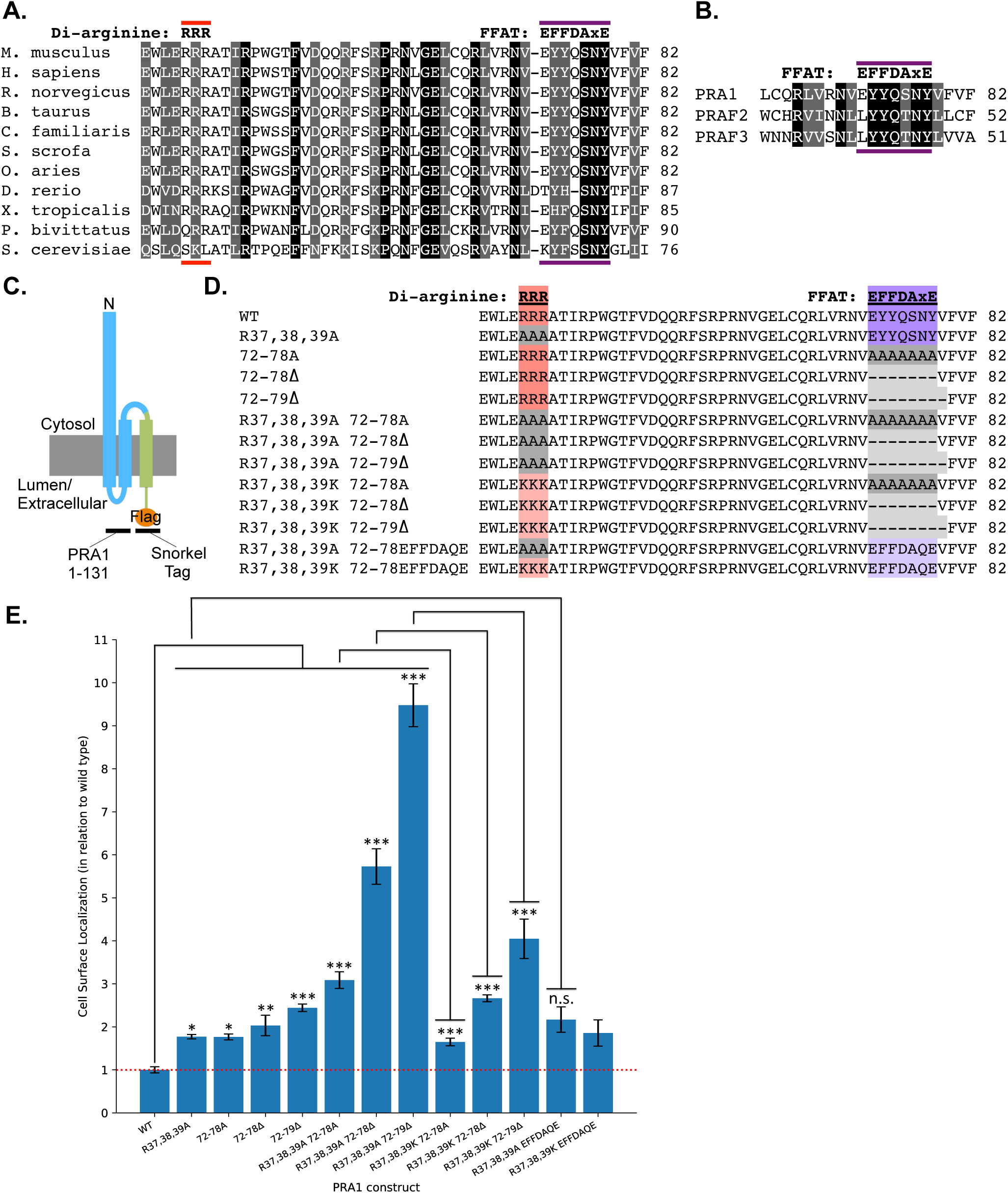
Targeted mutation of the PRA1 cytosolic N-terminal region reveals that di-arginine and FFAT-like motifs retain PRA1 constructs intracellularly. A cell surface assay carried out using COS-7 cells demonstrates that ER retention of a PRA1 construct with incomplete Golgi targeting information (amino acids 1-131) is driven by both di-arginine and FFAT-like motifs. (A) Alignment of PRA1 homologs shows that the di-arginine motif and the FFAT-like motif double aromatic residue signature are conserved in higher eukaryotes. (B) Alignment of all three mammalian prenylated rab acceptor family members shows that the FFAT-like motif is conserved. (C) The PRA1^1-131^-Snorkel-Tag construct used to identify ER retention motifs on the cytosolic N-terminal region. (D) PRA1 amino acid sequences for constructs designed and used in the cell surface assay. (E) Cell surface assay shows that targeted mutation of di-arginine and FFAT-like motifs leads to a statistically significant increase in cell surface localization. Substitution of the positively charged lysine for the di-arginine motif or of the classic FFAT motif for the endogenous FFAT-like motif rescues the cell surface localization phenotype. (n=9 for all constructs). One-way ANOVA was completed, followed by an LSD post-hoc test. *p< .05, **p< .01, ***p< .001, n.s. = not significant

Examination of the PRA1 cytosolic N-terminal amino acid sequence shows that there is one potential di-arginine ER retention motif that is both highly conserved, and lies within a functional distance from the membrane (Figure 3A) (Schutze et al., 1994; Shikano and Li, 2003). Upon mutation of the sequence to alanine (R37,38,39A), we observed an increase in cell surface localization (Figure 3D-E). After further examination of the N-terminal amino acid sequence, we identified a highly conserved string of residues that comprise a potential FFAT-like motif preceding the first predicted membrane insertion site (Fig. 3A). Although the FFAT motif was initially described as two phenylalanines (<u>FF</u>) in an <u>A</u>cidic <u>T</u>ract, strong deviations from this sequence have been identified (Loewen et al., 2003; Mikitova and Levine, 2012; Murphy and Levine, 2016). The initial FFAT motif consensus sequence was posited to be EFFDAxE, where x can be any amino acid. Subsequently, other aromatic residues have been found to substitute for phenylalanines and it has also been demonstrated that the acidic tract does not need to be completely intact (Murphy and Levine, 2016). The putative PRA1 FFAT-like motif contains an initial acidic amino acid (E), two aromatic residues (YY), a polar amino acid (Q), and SNY (Fig. 3A). Amino acid alignment shows that this motif is conserved in lower organisms including the budding yeast (Fig. 3A). Mutation of the seven amino acids that comprise the FFAT-like motif to alanine (72-78A) led to an increase in cell surface localization (Fig. 3D-E). Combining this mutation with the di-arginine to alanine mutant led to a strong and additive effect on the cell surface localization of the resulting construct (R37,38,39A/72-78A) (Fig. 3D-E). We found that deletion of the FFAT-like motif is much more effective at abrogating ER retention (72-78Δ and R37,38,39A/72-78Δ) (Fig. 3D-E). Amino acid alignment of PRA1 with the two other mammalian prenylated rab acceptor family members PRAF2 (JM4) and PRAF3 (Arl6ip5/Gtrap3-18) shows that the FFAT-like motif lies within the most conserved region (Fig. 3B). The alignment also shows that there is a conserved hydrophobic amino acid immediately downstream of the FFAT-like motif. Deletion of this amino acid along with the FFAT-like motif led to the largest increase in cell surface localization (72-79Δ and R37,38,39A/72-79Δ suggesting it plays an important role in this variant of the classic FFAT sequence (Fig. 3D-E).

It has previously been found that di-lysine and di-arginine motifs are often interchangeable, as both are used as positive charges that recruit the COPI coat, which facilitates ER retrieval (Beck et al., 2009; Zerangue et al., 2001). We replaced the alanines within the di-arginine/FFAT double mutants with lysines [(R37,38,39K/72-78A), (R37,38,39K/72-78Δ), and (R37,38,39K/72-79Δ)] and found that re-introduction of the positive charge leads to a significant decrease in cell surface localization compared to di-arginine/FFAT double mutants (Fig. 3D-E). We also found that substitution of one of the originally identified classic ORP3 FFAT motifs leads to a strong rescue of the cell surface localization phenotype (R37,38,39A/EFFDAQE) (Fig. 3D-E) (Loewen et al., 2003). Comparison of the di-arginine mutant with an intact native PRA1 FFAT-like motif (R37,38,39A) and the rescue (R37,38,39A/EFFDAQE) shows that the difference in cell surface localization between these two constructs is not statistically significant, suggesting that the novel FFAT-like motif described here is as effective at ER retention as the classic FFAT motif within the context of the PRA1 amino acid sequence (Fig. 3D-E).

Together, this data strongly suggests that the PRA1 cytosolic N-terminal region contains both a membrane distal di-arginine ER retrieval motif and a membrane proximal FFAT-like motif that mediate ER retention.

### PRA1 constructs with incomplete Golgi targeting information are retained within the ER and secreted to the cell surface upon mutation of both di-arginine and FFAT-like motifs

We sought to confirm the cell surface assay data by localizing constructs fused to the Snorkel-Tag within the cell via immunocytochemistry. Expression of the wild type PRA1^1-131^ construct that lacks complete Golgi targeting information in COS-7 cells leads to a reticular ER localization phenotype (Fig. 4A). Mutation of the di-arginine motif variant uncovered by the cell surface assay (R37,38,39A) led to a disruption of this reticular pattern, displaying a more particulate appearance (Fig. 4B). Deletion of the FFAT-like motif (72-78Δ) also led to a deviation from the wild type localization pattern, displaying a dispersed distribution (Fig. 4C). Double mutation of both of these ER retention/retrieval sequences (R37,38,39A/72-78Δ the loss of much of the reticular pattern (Fig. 4D). This construct is distributed diffusely throughout the cell with only some peripheral and disorganized reticular ER localization observed. Together, these results further corroborate the cell surface assay data, indicating that the PRA1 di-arginine and FFAT-like motifs within the cytosolic N-terminal region are necessary for proper ER retrieval/retention.

**Fig. 4.**
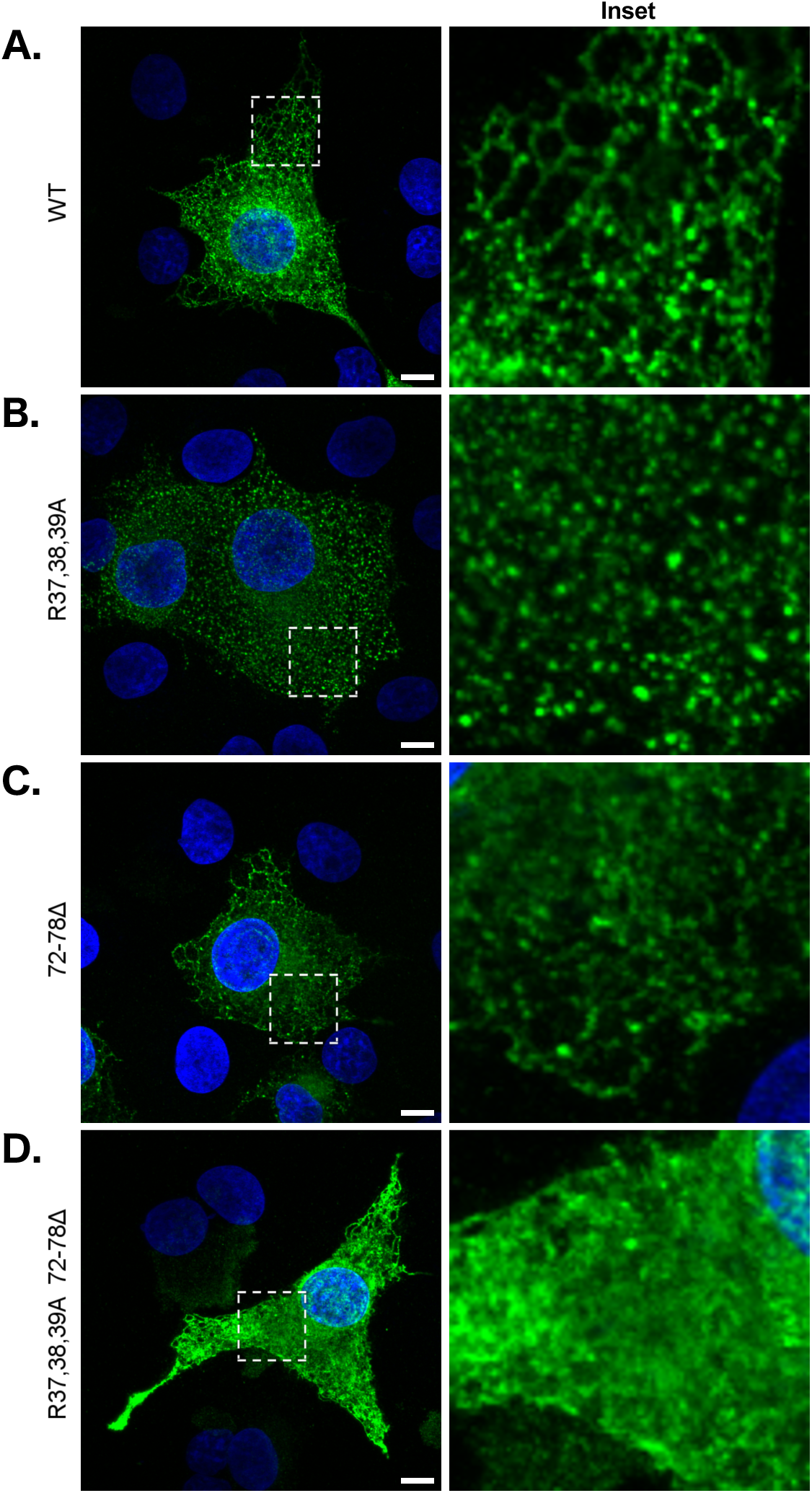
PRA1 constructs with incomplete Golgi targeting information are retained within the ER and secreted to the cell surface upon mutation of both di-arginine and FFAT-like motifs. The PRA1^1-131^ truncation construct, previously demonstrated to be retained within the ER, was fused to the Snorkel-Tag. The following constructs were expressed in COS-7 cells: (A) WT, (B) R37,38,39A, (C) 72-78Δ, 749 and (D) R37,38,39A/72-78Δ. Constructs were immuno-localized using an anti-FLAG antibody. Images are representative of at least three transfections and are compiled from a 6 μm z-section. Insets are magnifications of an area within the displayed images, highlighted with a dashed-line. Scale bar: 10 μm. Green: PRA1, Blue: nuclear labeling via DAPI

### Full length PRA1 is differentially localized when tags are added to either the N- or C-terminus

Previously published data describing PRA1 localization in mammalian cells has illustrated that PRA1 is a Golgi resident. Many of these studies relied on the expression of N-terminal tagged exogenous constructs (Abdul-Ghani et al., 2001; Gougeon et al., 2002; Hutt et al., 2000; Jung et al., 2011; Liang and Li, 2000; Liu et al., 2006). We have confirmed these observations by co-expressing GFP-PRA1 and the Golgi marker MannII-mCherry (Fig. 5A). In order to determine whether addition of GFP to the PRA1 C-terminus leads to a similar localization phenotype, we expressed PRA1-GFP and found that this construct displays a reticular localization pattern that strongly co-localizes with the mCherry-Sec61b ER marker (Fig. 5B).

**Fig. 5.**
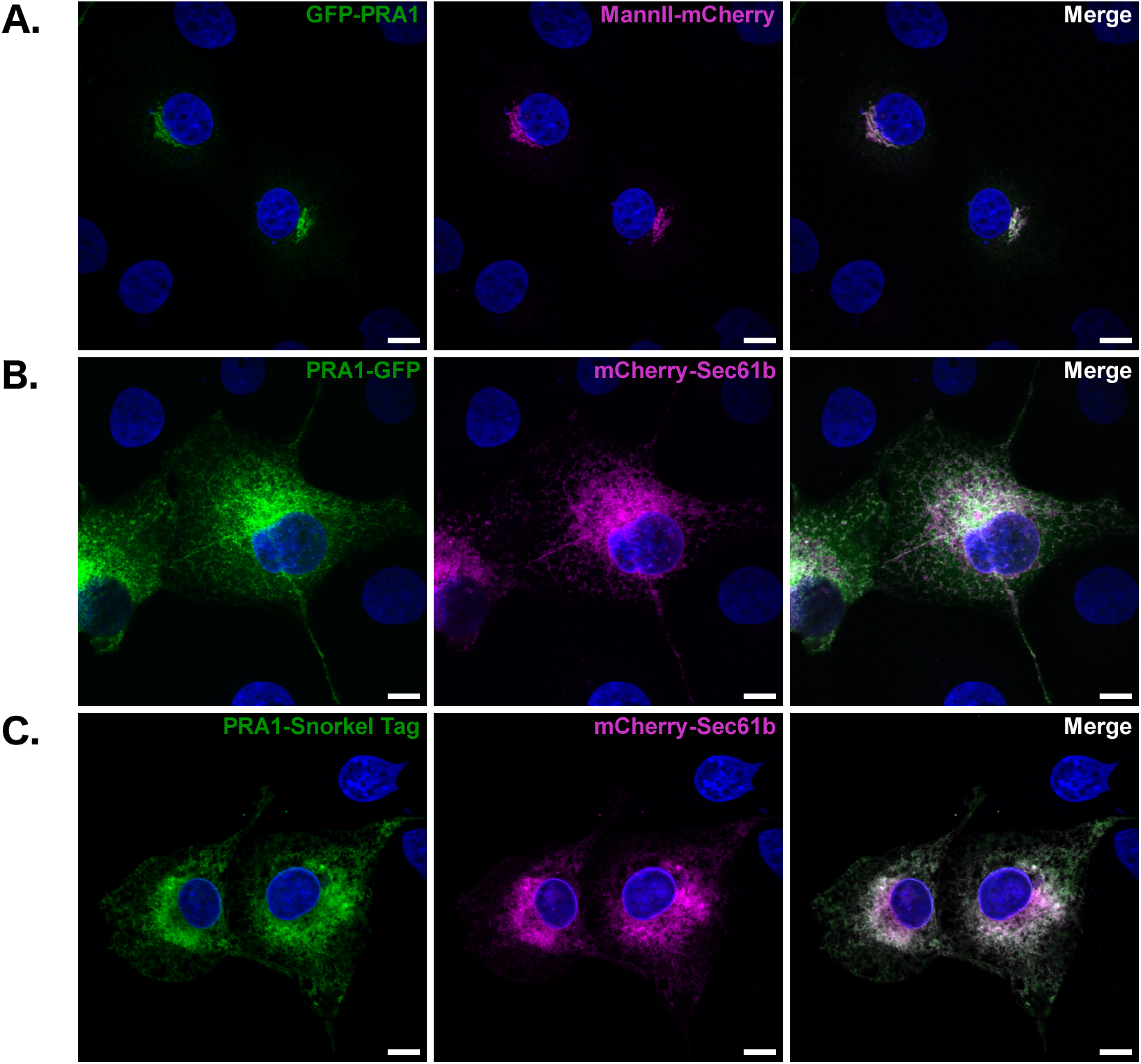
Full length PRA1 is differentially localized when tags are added to either the N- or C-terminus. GFP fusion to the PRA1 N-terminus leads to Golgi localization and GFP fusion to the C-terminus leads to reticular ER localization. The following constructs were expressed in COS-7 cells: (A) GFP-PRA1/MannII-mCherry (B) PRA1-GFP/mCherry-sec61b (C) PRA1-Snorkel-Tag/mCherry-sec61b. Images are representative of at least three transfections and are compiled from a 2 μm z-section. Scale bar: 10 μm. Green: PRA1 fusion constructs, Magenta: organelle marker label, Blue: nuclear labeling via DAPI, White: co-localization

Others have reported that the addition of the smaller HA tag to the N-terminus leads to the same phenotype seen here for the GFP-PRA1 construct (Gougeon et al., 2002; Hutt et al., 2000; Jung et al., 2011; Liu et al., 2006). This suggests that the size of the GFP tag is not the source of the differential trafficking observed here. Addition of the Snorkel-Tag to the C-terminus of PRA1 produced the same reticular pattern observed for PRA1-GFP constructs (Fig. 5C). Since the Snorkel-Tag is designed to have an epitope within the lumen, this suggests that a C-terminal cytosolic tag is not what drives reticular ER localization. Furthermore, we added a FLAG tag within the central cytosolic loop (PRA1^1-118-FLAG-119-185^) and between the first (PRA1^1-93-FLAG-94-185^) and second (PRA1^1-144-FLAG-145-185^) pair of predicted membrane spanning alpha-helices, but found that the resulting constructs were either localized to the cytosol or aggregated in a punctate manner, suggesting errors in folding (data not shown). To address the possibility of steric hindrance, we also added both helical and flexible linkers of up to 20 amino acids in length between GFP and the PRA1 N/C-termini and did not observe a significant deviation from the localization phenotype of constructs without linkers (data not shown) (Arai et al., 2001).

This data demonstrates that full length PRA1 localization is particularly sensitive to the addition of tags. An N-terminal tag renders ER retention motifs on the cytosolic N-terminus ineffective, while addition of a tag to the C-terminus results in a reticular ER localization phenotype. Neither of these constructs truly mirrors endogenous protein localization in both the retina and NIH3T3 cells (Figs. 1-2, S2). Together, these results highlight the importance of a cautious approach to the interpretation of full-length PRA1 localization data that is dependent on the expression of covalently tagged constructs.

## DISCUSSION

In the *rd1* mouse model of retinitis pigmentosa, photoreceptors degenerate during maturation, prior to the development of the outer segment. To better understand the degenerative process, we examined changes in gene expression prior to observable cell death at postnatal day 10. We found that PRA1 is significantly down-regulated in the *rd1* retina prior to the onset of degeneration (Dickison et al., 2012). PRA1 had been identified as a ubiquitously expressed Golgi resident and characterized as a regulator of Rab GTPase trafficking. Using a novel anti-PRA1 antibody, we confirm previous localization results and expand the cellular territory on which endogenous PRA1 resides within the secretory pathway in photoreceptors and cultured epithelial cells. To our knowledge, this is the first time that endogenous PRA1 has been localized to the ER in mammalian cells and the first report that demonstrates that endogenous PRA1 is enriched at ER-mitochondria membrane contact sites in any cell type. The latter observation has important functional implications.

Previous experiments have demonstrated that deletion of the PRA1 cytosolic C-terminal region, which contains Golgi targeting information, leads to extensive co-localization of the construct with Calnexin, a well characterized ER resident (Hutt et al., 2000). Consistent with this result, the data presented here suggest that abrogation of any amino acid sequence information known to be required for Golgi delivery leads to ER retention by default. Golgi targeting information has previously been found to reside on the PRA1 cytosolic C-terminal region (Abdul-Ghani et al., 2001; Liang et al., 2004). Specifically, the C-terminal domain contains a di-acidic motif, DGE, which is required for ER exit; PRA1 has been characterized as a member of COPII vesicles, suggesting that Golgi delivery is facilitated by the COPII coat complex (Abdul-Ghani et al., 2001; Jung et al., 2011; Otte et al., 2001). We interrogated the PRA1 cytosolic N-terminal region and found that two previously unidentified trafficking motifs facilitate ER retention, a membrane distal di-arginine motif and a membrane proximal FFAT-like motif.

The di-arginine motif is known to facilitate ER localization by recruiting the COPI coat (Yuan et al., 2003). We have previously completed an unbiased split-ubiquitin yeast two-hybrid screen to identify novel PRA1 binding partners and better understand its function in-vivo (Abu Irqeba and Ogilvie, 2019). Among the new binding partners we uncovered and confirmed in mammalian 1-COP, a member of the heptameric COPI coat (Kuge et al., 1993). This result is cells was ζ consistent with a report that identified PRA1 to be among the proteins that reside within purified COPI vesicles (Gilchrist et al., 2006). Together with our results, the data supports a model in which the COPI coat machinery plays an important role in PRA1 retrieval to the ER.

Others have localized PRA1 to the ER in both yeast and plants, but it is important to note that these studies relied on C-terminally tagged PRA1 constructs (Alvim Kamei et al., 2008; Geng et al., 2005). We found that although the di-arginine motif is not conserved in homologs within yeast and *Arabidopsis*, the FFAT-like motif is (Fig. 6A), suggesting that the ER localization that has been previously observed in both these organisms is mediated by this motif. Sequence alignment shows that the FFAT-like motif, but not the di-arginine motif, is also conserved in both PRAF2 and PRAF3 (Fig. 3B), which have been found to be ER residents (Doly et al., 2016; Ruggiero et al., 2008). This suggests that the FFAT-like motif described here drives ER retention of all three members of the prenylated rab acceptor family. With the motifs identified in this study, we update the model that describes PRA1 trafficking (Fig. 6B). The amino acid sequences that facilitate ER exit and Golgi delivery are on the PRA1 C-terminal region and the motifs that retain PRA1 within the ER are on the cytosolic N-terminal region.

**Fig. 6.**
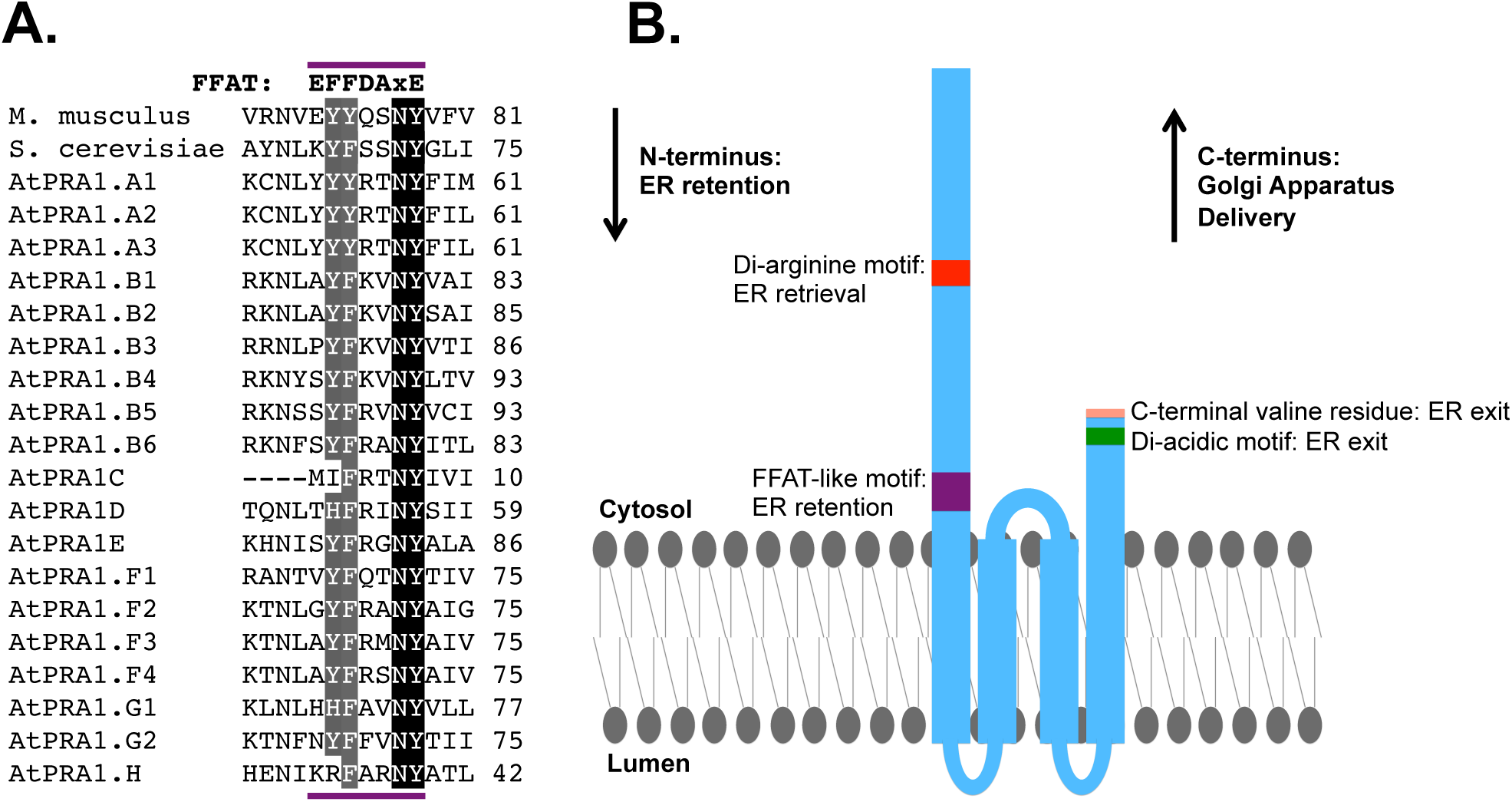
Proposed model for PRA1 trafficking in mammalian cells. (A) Sequence alignment of mouse, yeast, and *Arabidopsis* PRA1 family members shows that the FFAT motif consensus double aromatic residue signature in mammals is also conserved in both fungi and plants. The amino acids NY, which correspond to the “xE” in the classic FFAT motif are completely conserved in mammals, yeast, and *Arabidopsis* family members. (B) Schematic map of the trafficking motifs superimposed onto full length PRA1. The cytosolic C-terminal region contains both a C-terminal valine and a di-acidic motif previously demonstrated to facilitate ER exit.

Using an antibody specific for the N-terminal region, we found that the largest subpopulation of PRA1 resides at ER-mitochondria membrane contact sites in NIH3T3 cells (Fig. 2). This observation was confirmed with a polyclonal antibody generated against the full-length protein (Fig. S2). Our data further demonstrates that endogenous PRA1 localization does not mirror that of full-length, tagged PRA1 constructs expressed in mammalian cells (Fig. 5). Previous work showing that PRA1 is only enriched in the Golgi appears to rely on either N-terminally tagged constructs, or on antibodies with specificity for an epitope that may only be accessible at this organelle (Liang and Li, 2000; Liu et al., 2006; Liu et al., 2011). Notably, a phenotypic change in localization for tagged constructs has also been observed for Gp78/AMFR, another protein that lies at ER-mitochondria membrane contact sites (Registre et al., 2004). Large-scale studies in yeast have also led to observed changes in localization for many proteins depending on whether a tag is added to the N- or C-terminus (Weill et al., 2019; Weill et al., 2018).

Although previous studies have associated PRA1 with Rab GTPase trafficking within the secretory pathway, the first established links were based on yeast two-hybrid screens that did not take into account the fact that PRA1 is a transmembrane protein (Bucci et al., 1999; Calero and Collins, 2002; Janoueix-Lerosey et al., 1995; Martincic et al., 1997). Many of the studies that suggest PRA1 is a regulator of Rab GTPase trafficking are highly reliant on in-vitro data (Hutt et al., 2000; Ohya et al., 2009; Sivars et al., 2003). Subsequent in-vivo studies in both yeast and mammalian cells failed to produce any changes in wild type Rab GTPase localization after knock-out/knock-down of PRA1 (Cabrera and Ungermann, 2013; Geng et al., 2005; Voss et al., 2019). PRA1 association with Rab GTPases appears remarkably promiscuous and extends to other prenylated proteins, including other members of the Ras superfamily (Bucci et al., 1999; Calero and Collins, 2002; Figueroa et al., 2001). The simple addition of a CAAX box to the C-terminus of free GFP leads to its interaction with PRA1, suggesting that the link between many lipidated proteins and PRA1 may hinge on the lipid tail itself rather than the primary amino acid sequence of potential binding partners (Figueroa et al., 2001). Pulldown of PRA1 from both mammalian cells and yeast leads to co-immunoprecipitation of mostly ER localized proteins and fails to show that PRA1 associates with Rab GTPases (Geng et al., 2005; Huttlin et al., 2015). Our split-ubiquitin yeast two-hybrid screen also failed to show that PRA1 interacts with Rab GTPases (Abu Irqeba and Ogilvie, 2019).

Overexpression of PRA1 inhibits the anterograde movement of proteins through the secretory pathway in both mammalian and plant cells (Gougeon et al., 2002; Lee et al., 2011). Overexpression of the PRA1 homolog in yeast leads to the expansion of the ER (Geng et al., 2005). In a report describing a screen for proteins that regulate protein trafficking from the ER to the Golgi, knock-down of PRA1 was found to affect both COPI and COPII coat staining in HeLa cells (Simpson et al., 2012). Knock-down of PRA1 leads to an increase in intracellular LDLR, which facilities cholesterol uptake (Pietiainen et al., 2013), consistent with the observation that knock-down in NPC cells leads to intracellular cholesterol accumulation (Liu et al., 2011). Many proteins that contain a FFAT motif have been linked to lipid transport (Wong et al., 2019). Lipid transfer proteins that contain FFAT motifs have been found to tether at ER membrane contact sites with the plasma membrane, peroxisomes, Golgi, and endosomes (Levine, 2004). Lipids are also transferred between the ER and mitochondria, but the identification of candidate proteins that facilitate lipid transfer at these contact sites has proven to be challenging (Wu et al., 2018). Our finding that PRA1 resides at ER-mitochondria membrane contact sites and has a highly conserved FFAT-like motif supports previous work suggesting that it may play a role in lipid homeostasis.

The lack of in-vivo evidence that PRA1 functions as a regulator of Rab GTPase trafficking, its promiscuous association with proteins that are lipidated, its localization to the ER-mitochondria membrane contact sites, and our identification of a FFAT-like motif on its cytosolic N-terminal region support the hypothesis that PRA1 has a role in lipid trafficking and/or metabolism. Further studies are needed to identify and characterize lipid-binding capabilities and to screen for changes in Golgi structure, ER-mitochondria membrane contact site dynamics, and in lipid localization within the early secretory pathway following PRA1 knockdown. Our work, combined with the observation that PRA1 knockdown results in deviations in cholestesterol sequestration (Liu et al., 2011), suggest that future studies on the propensity of PRA1 to bind cholesterol, its derivatives, and both upstream and downstream byproducts may provide critical insights into its function. Our work also provides a foundation for future studies exploring a possible role associated with lipid homeostasis for the other mammalian prenylated rab acceptor family members. Whether the previously documented interactions between PRA1 and lipidated proteins have any functional relevance in-vivo requires further study.

## ACKNOWLEDGEMENTS

We would like to thank Angela Marie Richmond, Virginia Dickison, Ju Zhang, Jiyao Zhu, Rebecca Girresch, Cynthia Montana, Yuqi Wang, Susan Spencer, Brian Downes, Jonathan Fisher, Amanda Eccardt, Joseph Corbo, Grant Kolar, David Ford, Meribeth Broadway, Dana Baum, and Shiming Chen for helpful comments, suggestions, training, lab space, and reagents. We would also like to thank Jack Kennell for his support and the Saint Louis University Department of Biomedical Engineering for lab space accommodations.

## Funding

This work was supported by the Eunice Kennedy Shriver National Institute of Child Health and Human Development [grant number HD064269 to JMO], Sigma Xi Grants-in-Aid of Research [grant number G20101015154863 to AA], and Saint Louis University Beaumont Faculty Development and Presidential Research Funds [to JMO]. AA was further supported by a fellowship from the Saint Louis University College of Arts and Sciences. Biology Department core facilities at Saint Louis University are supported by an equipment grant from the National Science Foundation [grant number DBI-0421383]. Some images were taken using equipment at the St. Louis University School of Medicine Research Microscopy and Histology Core.

## Competing Interests

Disclosure: **A. Abu Irqeba**, None; **J.M. Ogilvie**, None

## Author Contributions

AA conceived of the project, designed and executed experiments, collected and analyzed data, and wrote the manuscript. JMO supervised the project and wrote the manuscript.

