## Supplementary Methods and Table 1 for "Di-arginine and FFAT-like motifs retain a subpopulation of PRA1 at ER-mitochondria membrane contact sites"

| | |
|---|---|
| MluI_PRA1_1+FWD | GCTGCG**ACGCGT**GCGGCCCAGAAGGACCAGCAG |
| BglII_Stop_PRA1-185_RVRS | GCTGCG**AGATCT**TTACACAGGTTCCATCTGCAGCTCCTC |
| XhoI_PRA1_1+_FWD | GCTGCG**CTCGAG**ATGGCGGCCCAGAAGGACCAGCAG |
| MscI_PRA1_185+_RVRS | GATACAC**TGGCCA**CCACAGGTTCCATCTGCAGCTC |
| XbaI_XhoI_ATG_EGFP_FWD | GCTGCG**TCTAGA**CTCGAG**ATG**GTGAGCAAGGGCGAGGAG |
| BglII_STOP_MluI_EGFP_RVRS | GCTGCG**AGATCT**TCATCA**ACGCGT**CTTGTACAGCTCGTCCATGCC |
| mCHERRY_FWD_5'Cloning | GCTGCG**GAATTC**GACGCT**ACGCGT**GTGAGCAAGGGCGAGGAGGATAAC |
| mCHERRY_RVRS_5'Cloning | GCTGCG**AGATCT**TTA**CTTGTCATCGTCGTCCTTGTAGTC**CTTGTACAGCTCGTCCATGCCGCC |
| mCHERRY_FWD_3'Cloning | GCTGCG**GAATTC**CGCCACC**ATGGACTACAAGGACGACGATGACAAG**GTGAGCAAGGGCGAGGAGGATAAC |
| mCHERRY_RVRS_3'Cloning | GCTGCG**AGATCT**TTACGT**ACGCGT**CTTA**GGTACC**CTTGTACAGCTCGTCCATGCCGCC |
| KpnI_mSec61b_FWD | GCTGCG**GGTACC**CCGGGTCCAACGCCCAGTGGCACC |
| KpnI_mSec61b_RVRS | GCTGCG**ACGCGT**TTATGATCGCGTGTACTTGCCCCAAAT |
| XbaI_mMannII_FWD | GCTGCG**TCTAGA**ATGAAGTTAAGTCGCCAGTTCACCGTG |
| Mlu_mMannII-aa116_RVRS | GCTGCG**ACGCGT**CAAACAGTCTCTGGGGTCAGCCTG |
| KpnI_ PRA1_131+RVRS | GCTGCG**GGTACC**CTGATGTGCTGGGCTCACCTC |
| PRA1_R37/38/39/43A_FWD | **GAGGCCGCCGCCGCGACCATCGCCCCCTGG** |

| | |
|---|---|
| PRA1_R37/38/39/43A_RVRS | GGCGATGGTCGCGGCGGCGGCCTCCAG |
| PRA1_R37/38/39K_FWD | CTGGAGAAGAAGAAGGCGACCATCCGG |
| PRA1_R37/38/39K_RVRS | CCGGATGGTCGCCTTCTTCTTCTCCAG |
| PRA1_80-86_OE_FWD | TTCGTGTTTCTCGGCCTCATC |
| PRA1_72-78A_RVRS | **GCCGAGAAACACGAA**CACGGCGGCGGCGGCGGCGGCGGCCACGTT |
| mPRA1_66-86_72-78 Δ _RVRS | GATGAGGCCGAGAAACACGAACACCACGTTGCGTACCAGGCG |
| mPRA1_66-86_72-79 Δ _RVRS | GATGAGGCCGAGAAACACGAACACGTTGCGTACCAGGCG |
| mPRA1_YYQSNY-FFDAQE | GAGAAACACGAACAC**CTCCTGGGCGTCGAAGAA**CTCCAC |

**Table S1. Primer sequences used in this study**

### Supplementary Methods

*ShRNA design*

ShRNA constructs were made according to previously described protocols (Fellmann et al., 2013). Mir30 based 97nt hairpins were amplified using the following primers: 5'-cagaaggctcgagaaggtatattgctgttgacagtgagcg-3' and 5'-tctcgaattctagccccttgaagtccgaggcagtaggc-3' with the reverse primer containing the "CNNC" motif retrofitted for more efficient processing (Fellmann et al., 2013). The resulting PCR fragment was cut with XhoI/EcoRI and ligated into the pCAG-mir30 vector (Addgene plasmid #14758) (Matsuda and Cepko, 2007).

Contructs generated (5'-3'): hPRA1-a-mir30, [tgctgttgacagtgagcgCGCGCAGAAGGACCAGCAGAAAtagtgaagccacagatgtaTTTCTGCTGGTCCTTCTGCGCTtgcctactgcctcgga]

hPRA1-b-mir30,
[tgctgttgacagtgagcgAACCCTGCTGCCGAAGCTGATTtagtgaagccacagatgtaAATCAGCTTCGGCAGCAGGGTCtgcctactgcctcgga]
hPRA1-c-mir30,
[tgctgttgacagtgagcgCGGTGGCTCTGGCTGTCTTTTTtagtgaagccacagatgtaAAAAAGACAGCCAGAGCCACCAtgcctactgcctcgga]
GFP-mir30,
[tgctgttgacagtgagcg**aagccacaacgtctatatcatg**tagtgaagccacagatgta**catgatatagacgttgtggctg**tgcctactgcctcgga]. Prior to use, constructs were verified via Sanger sequencing at Eurofins Sequencing (Louisville, KY, USA).

*Untagged PRA1 construct*

PRA1 was amplified from mouse cDNA purchased from Origene (see Materials and Methods). The amplified product was cloned into the PCAGIG vector (Addgene-see Materials and Methods) using the XhoI and BglII restriction enzyme sites.

*Generation of Stable ShRNA cell lines (Abgent anti-PRA1 antibody confirmation)*

To generate selectable ShRNA constructs, the generated mir30 constructs were amplified from the pCAG-mir30 backbone and subcloned within the pcDNA 3.1 (-) vector XbaI/BamHI restriction enzyme sites. Resulting constructs were sequence verified and linearized using the BglII restriction enzyme site. 1.5 µg of each construct was then transfected into Hek293T cells using Lipofectamine 2000 (12 well plates). 24 hrs later, cells were trypsinized and moved to 100 mm cell culture plates where selection was carried out over a 10 day period. An untransfected control plate showed complete cell death after the selection period was finished. Media was changed every 3 days with fresh Hygromycin B added each time.

**Fellmann, C., Hoffmann, T., Sridhar, V., Hopfgartner, B., Muhar, M., Roth, M., Lai, D. Y., Barbosa, I. A., Kwon, J. S., Guan, Y. et al.** (2013). An optimized microRNA backbone for effective single-copy RNAi. *Cell Rep* **5**, 1704-13.

**Matsuda, T. and Cepko, C. L.** (2007). Controlled expression of transgenes introduced by in vivo electroporation. *Proc Natl Acad Sci U S A* **104**, 1027-32.
