## Supplementary Figures 1-2 for "Di-arginine and FFAT-like motifs retain a subpopulation of PRA1 at ER-mitochondria membrane contact sites"

#### Supplementary Figure 1.

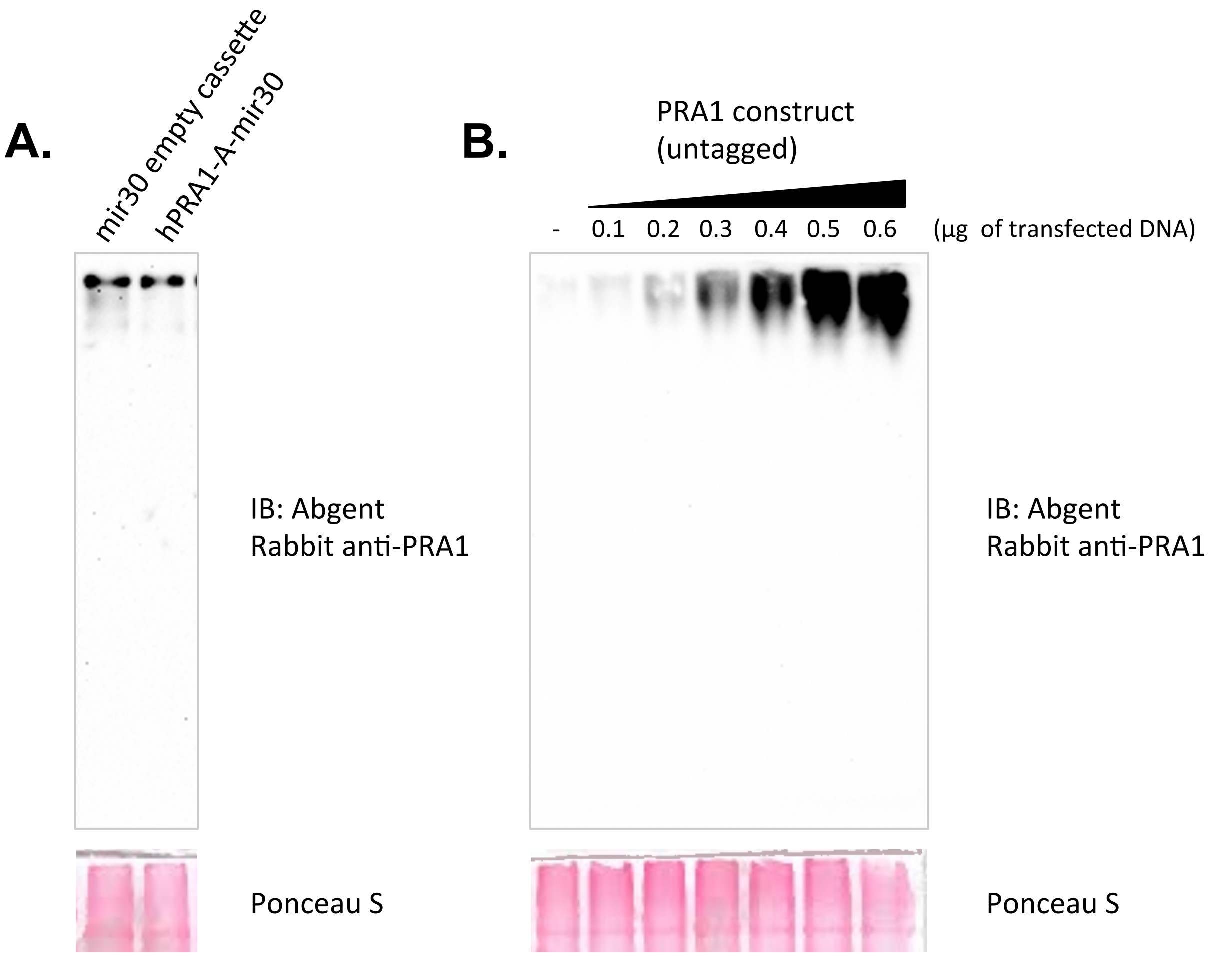


**Fig. S1. Confirmation of Abgent rabbit anti-PRA1 antibody (Cat #AP9049a) using native-PAGE western blots.**
(A) Stably transfected polyclonal Hek293T cell lines that express both an empty and hPRA1 targeting mir30 cassettes were generated. PRA1 knock-down leads to a depletion of bands detected by the Abgent anti-PRA1 antibody.
(B) An increasing amount of an untagged mouse PRA1 cDNA construct was expressed in Hek293T cells. The bands detected using the Abgent anti-PRA1 antibody correspond in size and increase in intensity in accordance with the increasing amount of construct delivered to Hek293T cells.

### Supplementary Figure 2.

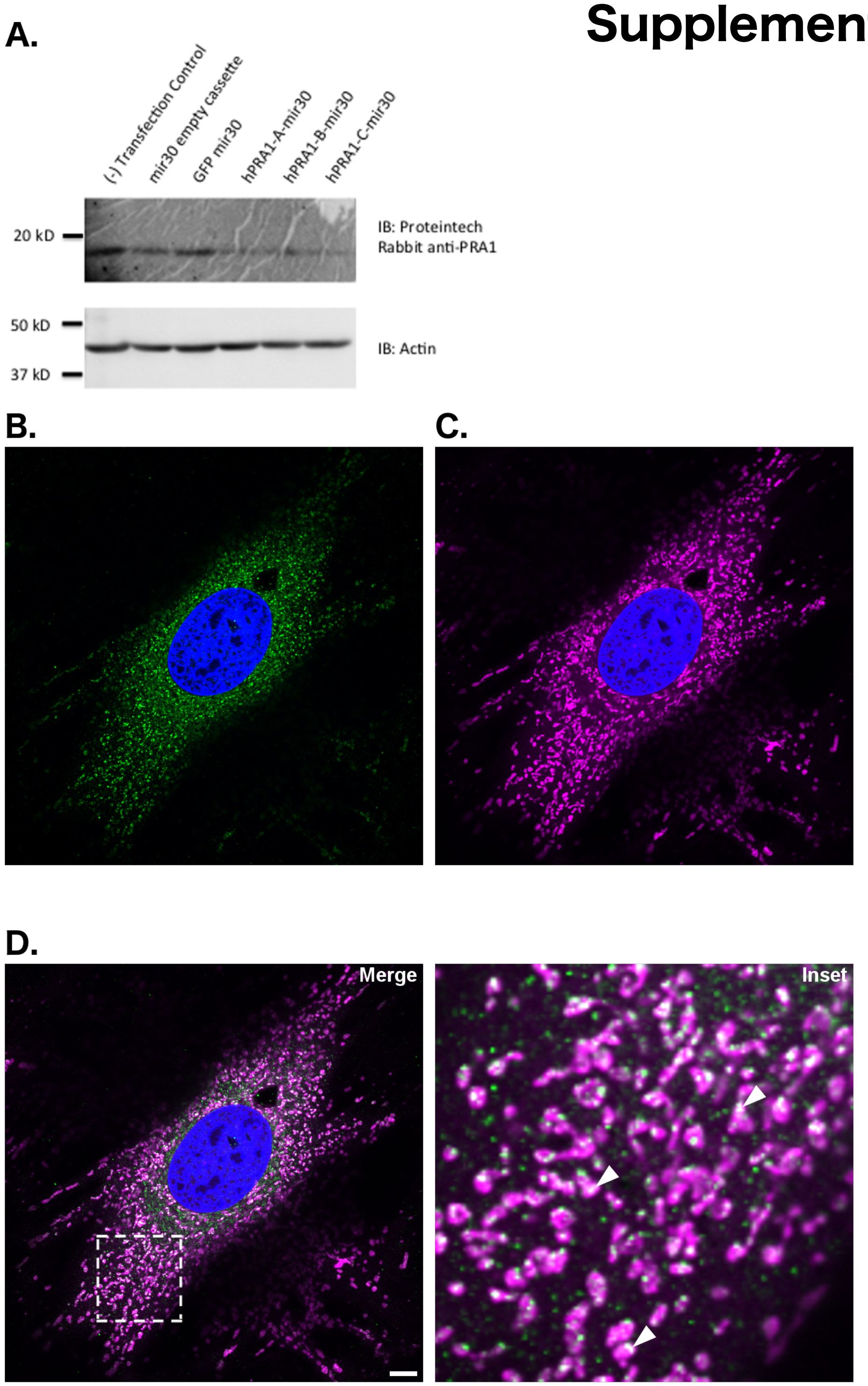


**Fig. S2. Proteintech anti-PRA1 antibody shows that a subpopulation of endogenous PRA1 co-localizes with Mitochondria in NIH3T3 cells.** (A.) A band is detected at the predicted size by the Proteintech rabbit anti-PRA1 antibody (Cat. #10542-1-AP) after standard western analysis. The detected band is depleted upon targeted knock-down in Hek293T cells. (B.) Immunohistochemical localization of PRA1 using the Proteintech anti-PRA1 antibody and the MitoTracker marker (C.) in NIH3T3 cells shows co-localization (see D.). Solid arrowheads denote regions of co-localization. Images are representative of at least three acquisitions and are compiled from a 1 μm z-section. The inset is a magnifications of an area within the displayed image, highlighted with a dashed-line. Scale bar: 10 μm. Green: PRA1, Magenta: Mitotracker, Blue: nuclear labeling via DAPI, white: colocalization
